# Designed fluorescent protein cages as fiducial markers for targeted cell imaging

**DOI:** 10.1101/2024.02.28.582585

**Authors:** Morgan Gee, Kaiser Atai, Hilary A. Coller, Todd O. Yeates, Roger Castells-Graells

## Abstract

Understanding how proteins function within their cellular environments is essential for cellular biology and biomedical research. However, current imaging techniques exhibit limitations, particularly in the study of small complexes and individual proteins within cells. Previously, protein cages have been employed as imaging scaffolds to study purified small proteins using cryo-electron microscopy (cryo-EM). Here we demonstrate an approach to deliver designed protein cages – endowed with fluorescence and targeted binding properties – into cells, thereby serving as fiducial markers for cellular imaging. We used protein cages with anti-GFP DARPin domains to target a mitochondrial protein (MFN1) expressed in mammalian cells, which was genetically fused to GFP. We demonstrate that the protein cages can penetrate cells, are directed to specific subcellular locations, and are detectable with confocal microscopy. This innovation represents a milestone in developing tools for in-depth cellular exploration, especially in conjunction with methods such as cryo-correlative light and electron microscopy (cryo-CLEM).

## INTRODUCTION

Visualizing the spatial and temporal organization of proteins inside cells is essential for understanding their biological functions and their role in cellular processes ^1^. *In vivo* imaging of proteins has played a crucial role in rapidly developing therapeutics, specifically in areas of research such as cancer biology and drug discovery ^2–5^. While fluorescence microscopy is a widely used technique for protein imaging, its application is limited by the need for specific protein labeling and by the limited resolution and photophysical properties of fluorescent proteins ^6,7^. Cryo-electron tomography (cryo-ET) offers higher resolution, but it is currently constrained to large assemblies due to the difficulty in identification of the molecule of interest in the crowded and complex environment of the cell ^8–10^.

To address these limitations, we have developed fluorescent designed protein cages that enable targeted imaging of proteins inside cells by acting as identifiable markers. Protein cages are hollow, nanoscale structures that can be designed to spontaneously self-assemble from multiple copies of modular protein subunits ^11–17^. The resulting cage structures have well-defined sizes and shapes and can be engineered to encapsulate guest molecules, display functional proteins on their surfaces, and work as imaging scaffolds for cryo-electron microscopy (cryo-EM) 18–26.

The protein cages presented in this study are genetically fused to a designed ankyrin repeat protein (DARPin) ^27^. Similar to antibody domains, DARPins have been exploited in recent work as a modular system for obtaining sequence variants that bind with high affinity to diverse proteins of interest; the DARPin in the present study binds to GFP ^28^. By using a DARPin as the targeting moiety, we can image proteins of interest with high specificity without the need for direct genetic manipulation of the target protein or the chemical labeling of cells. Protein cages also offer other prospective advantages over traditional protein imaging methods, including high stability, low cytotoxicity, and biocompatibility ^29^. They can be targeted to bind to any protein of interest, enabling imaging of previously inaccessible proteins. Additionally, our protein cages have a larger size compared to conventional fluorescent proteins, enabling improved localization accuracy. In addition to fluorescence microscopy, our designed protein cages show potential in applications in high-resolution cryo-correlative light and electron microscopy (cryo-CLEM) ^30^. The protein cages are designed to be large enough to be identifiable by the data processing software used to analyze tomograms obtained by electron microscopy, which could enable precise localization and 3D reconstruction of the target protein ^8,9,31^.

We describe the development and characterization of protein cages designed for targeted imaging of proteins inside cells **(Fig. 1)**. We demonstrate that our protein cages can be delivered into mammalian cells, can specifically bind to proteins of interest in the cytosol and in organelles, and can be imaged using confocal microscopy. Overall, the development of designed protein cages for intracellular delivery offers a promising approach for studying proteins in their native cellular context and for delivering cargo for a range of applications, including therapeutics, vaccine development, and catalysis ^23,25,32–39^.

**Figure 1.**
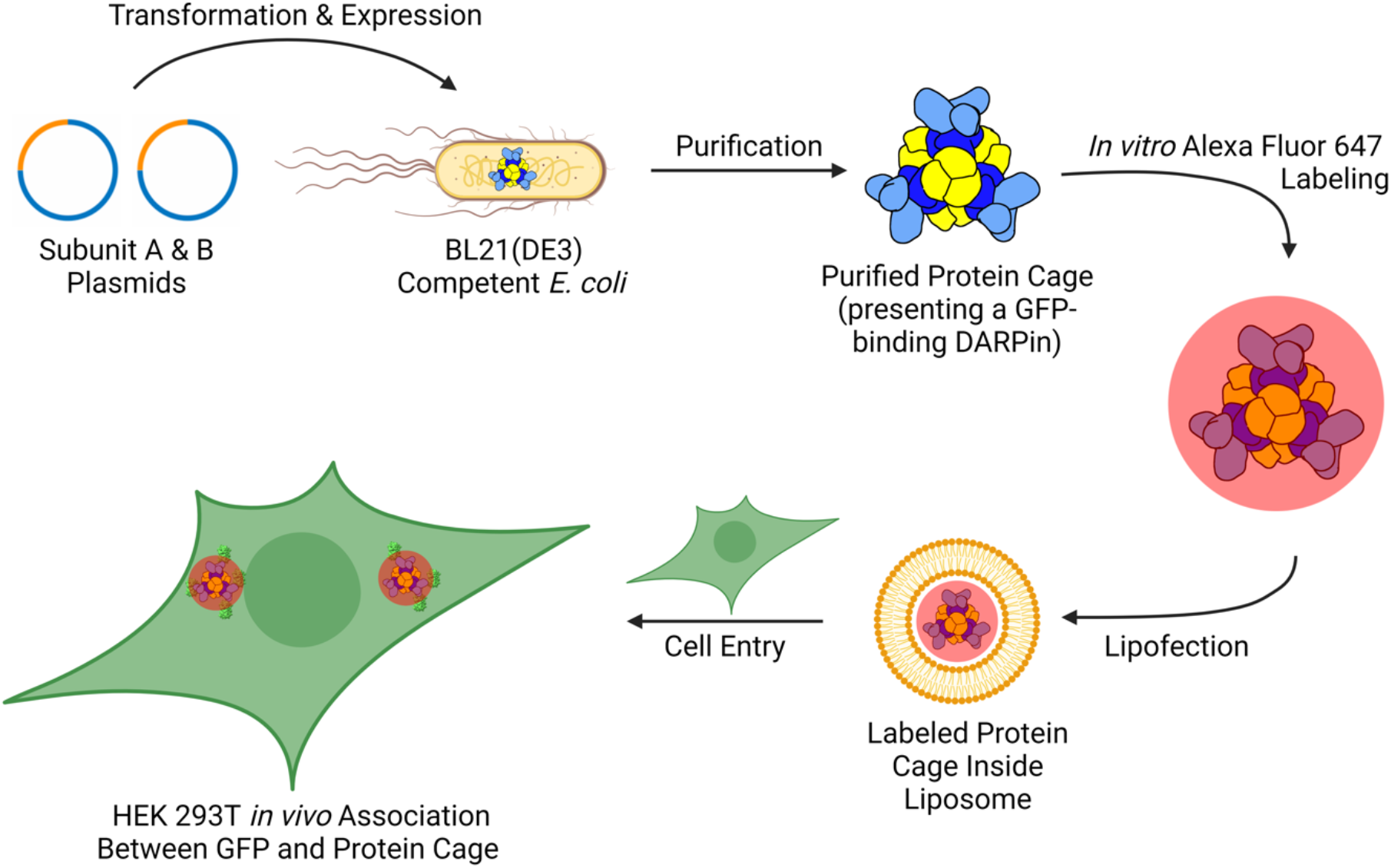
Workflow for the production, characterization, and lipofection of fluorescently labeled protein cages into HEK-293T cells. Plasmids containing the genes for the two subunits of the protein cage were transformed into *E. coli* cells followed by protein expression. The protein cages present a GFP-binding DARPin on their surface. Following the spontaneous, *in vivo* self-assembly of the protein cage, they were purified and fluorescently labeled with Alexa Fluor 647, a bright far-red fluorescent dye. These underwent lipofection to be transported into HEK-239T cells, integrating within liposomes as a payload for cell imaging. After an incubation period, colocalization of Alexa Fluor 647-labeled protein cages and endogenously expressed GFP within the HEK-239T cells occurred, which was visualized using confocal microscopy.

## RESULTS

### Characterization of Fluorescently Labeled Designed Protein Cages

As a basis for our nanoparticle imaging, we used a designed 12 nm tetrahedral protein cage (A12B12 subunit stoichiometry) with a genetically fused outward-facing DARPin that binds to green fluorescent protein (GFP), previously described in Castells-Graells *et al*., 2023. We transformed the A and B subunit genes of the protein cage into *Escherichia coli* BL21(DE3) cells. We purified the protein cages from cell lysate using Ni-NTA affinity chromatography based on a polyhistidine tail on the A subunit. To be able to subsequently label the protein cages with Alexa Fluor 647 (AF647), we had to modify existing protocols used for protein cage purification to avoid amine-based buffers, as the fluorophore needs to react with lysine residues of the protein to form covalent amide bonds (see Methods).

Following Ni-NTA affinity chromatography, we conducted size exclusion chromatography (SEC) and identified the presence of three differently sized protein populations, where the first peak represents the whole protein cage, the second peak represents trimer complexes, and the third peak represents the monomeric subunits **(Fig. 2a)**.

**Figure 2.**
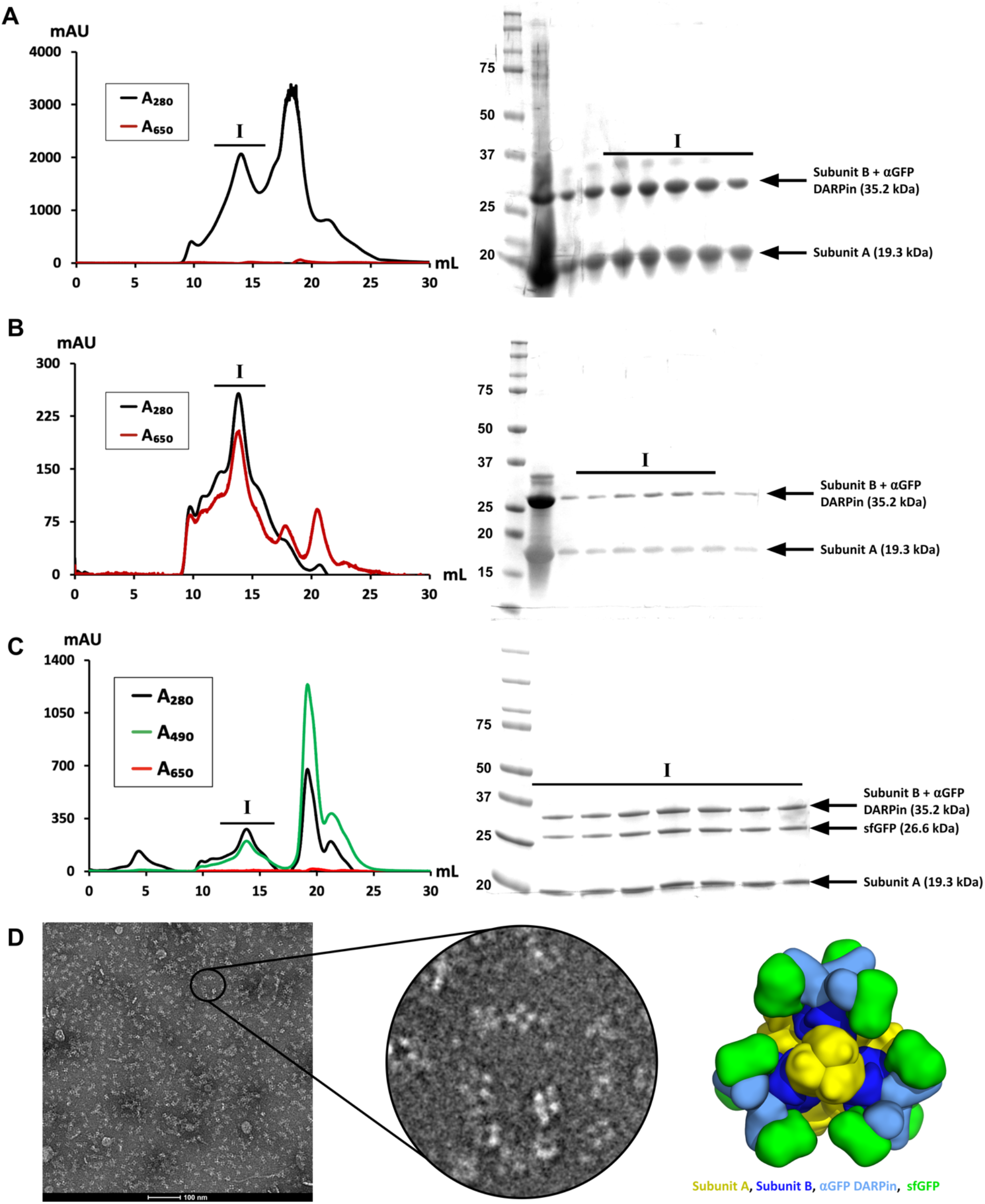
Characterization of fluorescently labeled protein cages. **(a)** SEC profile of unlabeled protein cage. Excitation at 650 nm was measured to determine naturally occurring absorbance that could interfere with detection of AF647. The SDS-PAGE band at 19.3 kDa supports the presence of subunit A, and the band at 35.2 kDa supports the presence of subunit B. **(b)** SEC profile of AF647-labeled cage. At peak I, there is significant absorbance at both 280 nm and 650 nm, representing the co-elution and the complex formation of assembled protein cage and AF647. The SDS-PAGE band at 19.3 kDa supports the presence of subunit A, and the band at 35.2 kDa supports the presence of subunit B. **(c)** SEC profile of sfGFP-bound cage. At peak I, there is significant absorbance at both 280 nm and 490 nm, representing the co-elution and the complex formation of assembled protein cage and sfGFP. The SDS-PAGE band at 19.3 kDa supports the presence of subunit A, the band at 26.6 kDa supports the presence of sfGFP, and the band at 35.2 kDa supports the presence of subunit B. **(d)** Negative stain electron microscopy images of AF647-labeled protein cages. Imaged properly assembled AF647-labeled protein cages could be visualized in a relatively monodisperse environment. An enhanced image of various protein cages can be compared to a 3D-reconstruction of a native protein cage.

The SEC plots demonstrate that there is little to no naturally occurring excitation at 650 nm, the absorbance maximum for AF647, exhibited by the protein cage subunits, allowing us to visualize successful labeling with fluorophore using both SEC and confocal microscopy **(Fig. 2a)**. SDS-PAGE analysis confirmed the presence of subunit A (19.3 kDa) and subunit B (35.2 kDa).

After the initial SEC purification, we labeled the purified protein cages with AF647 and performed another SEC purification to isolate the labeled complex **(Fig. 2b)**. We visualized the co-elution of the assembled protein cage and AF647 with two peaks at 280 nm and 650 nm, on the same fraction, for the protein cage and the fluorophore, respectively. This confirmed attachment of the AF647 and the protein cage. SDS-PAGE analysis confirmed the presence of the two distinct cage subunit types.

We also wanted to ensure that the assembled protein cages could properly associate with sfGFP, through binding to the genetically fused DARPin in the novel amine-free environment ^26,40^. After *in vitro* incubation of purified sfGFP and purified protein cage, we purified the sample with SEC to remove any disassembled cages and any unbound sfGFP **(Fig. 2c)**. sfGFP exhibits maximum absorption at 490 nm, allowing us to visualize complex formation between the sfGFP and assembled protein cage with co-elution of two peaks at 280 nm and 490 nm. Additionally, we noticed that there is no noticeable excitation at 650 nm in the absence of chemical fluorescent labeling, suggesting that there is no significant excitation caused by the complex formation of protein cage and sfGFP that could interfere with detection of AF647. SDS-PAGE analysis confirmed the presence of the two subunits from the protein cage as well as sfGFP (26.6 kDa).

Binding of sfGFP to the protein cages does not have a detrimental effect on their assembly ^19^, however it was not known if the covalent bonding of AF647 could have any significant deleterious effects. We used negative stain electron microscopy to visualize the purified AF647-labeled protein cages and confirm proper cage assembly in the presence of chemical labelling **(Fig. 2d)**.

### Internalization of Designed Fluorescent Protein Cages Into HEK-293T Cells

After the protein cages were successfully labeled with both AF647 and sfGFP, they were prepared for cellular experiments. For the first set, we wanted to determine if the protein cages could be properly internalized and, if so, whether the internalized protein cages were cytotoxic. To determine the capacity of protein cages to enter the cells, we used a lipofection-based approach using Lipofectamine CRISPRmax, which has been shown to effectively lipofect ribonucleoprotein complexes into mammalian cells ^41^. We predicted that lipofection would allow the cages to be efficiently packaged into liposomes and released into the cytoplasm. As an independent mode for tracking the cages, we used their DARPin domains to bind them to as many as twelve GFP molecules each *in vitro*, prior to incorporation into HEK-293T cells. With this approach, any green fluorescent puncta we observe intracellularly would be indicative of GFP-bound cages entering the cell. We tested a range of protein cage concentrations (from 2 µg/mL to 250 µg/mL) with incubation for 4 hours **(Fig. 3)**. After 4 hours of lipofection, we detected green fluorescent puncta in all the tested conditions, demonstrating that even at low concentrations the cages can be delivered into cells quickly and efficiently. As we increased the concentration of cages during lipofection, we found that there was a concomitant increase of intracellular green puncta, suggesting that the lipofection of these cages is tunable. Internalization of protein cages did not result in noticeable cytotoxicity.

**Figure 3.**
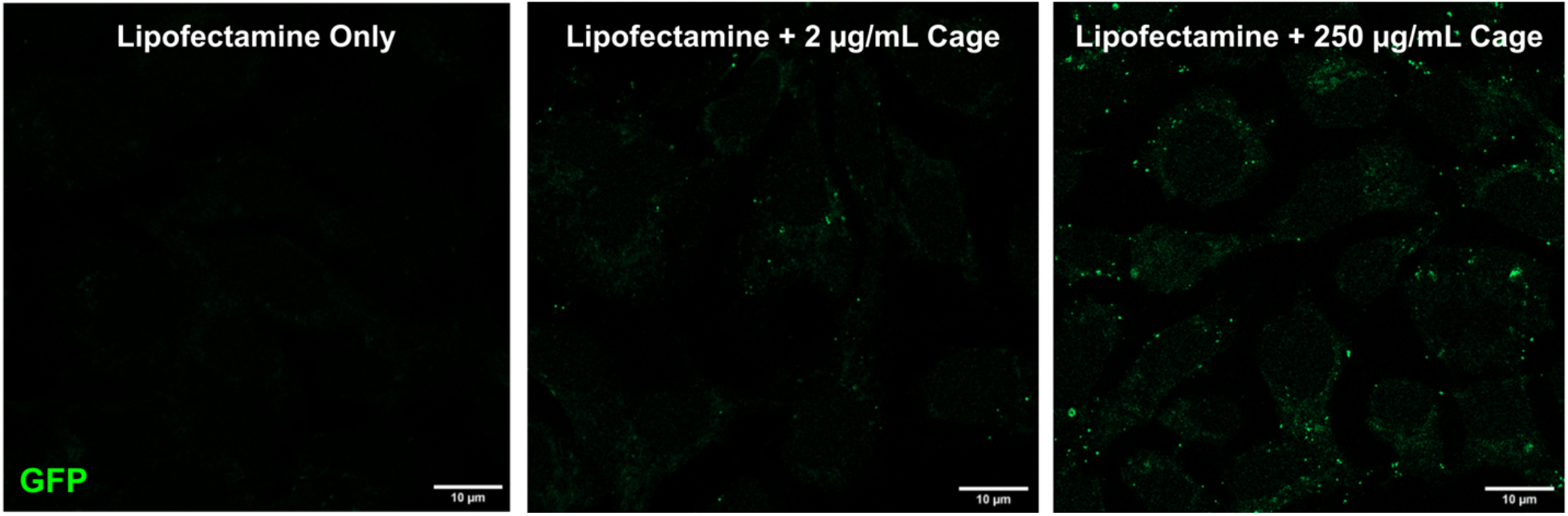
Visualization of green puncta as a result of internalized designed sfGFP-bound protein cages. After incubation of HEK-293T cells with varying concentrations of sfGFP-bound protein cages, green puncta can be visualized. For the condition with only Lipofectamine and no protein cages, there are no visible green puncta, which confirms that the laser/filter used does not detect any fluorescence without protein cages. As the concentration of the protein cages increases, so does the number of green puncta, suggesting that there is a positive correlation between protein cages and number of green puncta.

### Internalization of Designed Protein Cages into HEK-293T Cells and Binding to Endogenously Expressed GFP

After establishing that our designed protein cages can penetrate cells without any noticeable deleterious effects on the protein cages or on the cells, we wanted to determine if they are able to enter the cytoplasm. We theorized that if the cages are able to readily enter the cytoplasm, the exposed DARPins on the cages should nucleate any endogenously expressed GFP that is in the cytoplasm and produce bright green fluorescent puncta. Note, importantly, that each molecular cage binds (up to) 12 GFP proteins, making it feasible to detect individual particles (which would typically not be possible for a single fluorescent protein). Alternatively, if the protein cages are unable to enter the cytoplasm and instead remain inside the liposomes, we would not visualize green puncta. To test this, we used the same lipofection approach as before, but instead used cages that were not loaded with GFP and therefore had DARPins that were accessible and unbound. We then lipofected these “naked” cages into HEK-293T cells that express cytoplasmic GFP **(Fig. 4)**. After an incubation period, we were able to clearly visualize green puncta across all conditions with lipofected protein cages, while there was a clear absence of green puncta in the lipofectamine-only control. This demonstrates that even at low concentrations, protein cages can exit the liposome formed during the lipofection process and enter the cytoplasm to freely interact with any cytoplasmic protein to which they might be targeted (i.e. by their DARPin domain).

**Figure 4.**
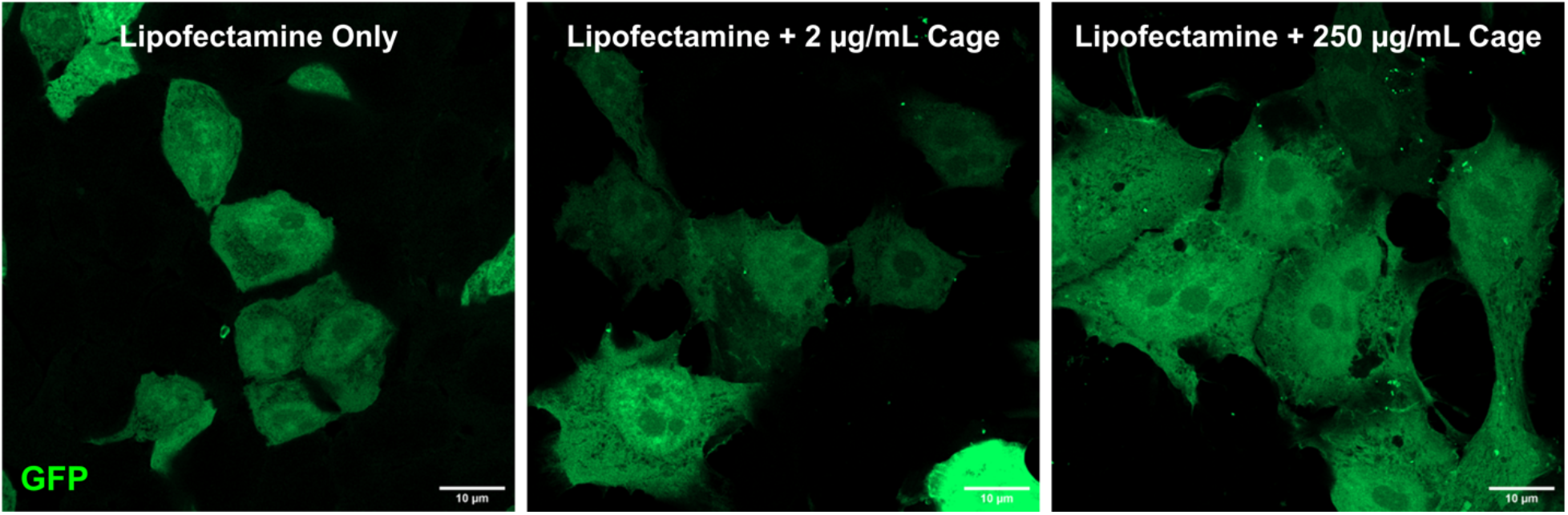
Visualization of green puncta formed by “naked” designed protein cages. After incubation of GFP-expressing HEK-293T cells with varying concentrations of “naked” protein cages, green puncta can be visualized. This suggests that the protein cages can escape the liposomes from the lipofection process and interact with cytoplasmic proteins.

### Internalization of Designed Alexa Fluor 647-labeled Protein Cages into HEK-293T Cells with Endogenous GFP

To confirm that the green puncta observed upon cage entry represent GFP bound to the cages, we performed colocalization experiments to visualize AF647-labeled protein cages and endogenously expressed GFP. Because AF647 has a maximum excitation of 650 nm and a maximum emission at 671 nm, we expect to see a clear difference in spectral emission compared to GFP. For these experiments, we followed the prior lipofection method but instead used cages that were fluorescently labeled with AF647. When transfecting the protein cages into HEK-293T cells expressing cytoplasmic GFP, we can clearly visualize AF647-positive puncta in the far-red spectral region, representing the fluorescent protein cages. We similarly see fluorescent puncta in the GFP-emission spectra, but only when cells are presented with the protein cages. Without the presence of the protein cages, the cytoplasmically expressed GFP and subsequent green fluorescent signal remains diffuse **(Fig. 5a-b)**. By merging the images, we can see the overlap of AF647 and green fluorescent puncta, demonstrating colocalization of the protein cages and endogenous GFP. This suggests that protein cages induce the formation of the green puncta by aggregating up to 12 endogenously expressed GFP molecules per protein cage **(Fig. 5b)**.

**Figure 5.**
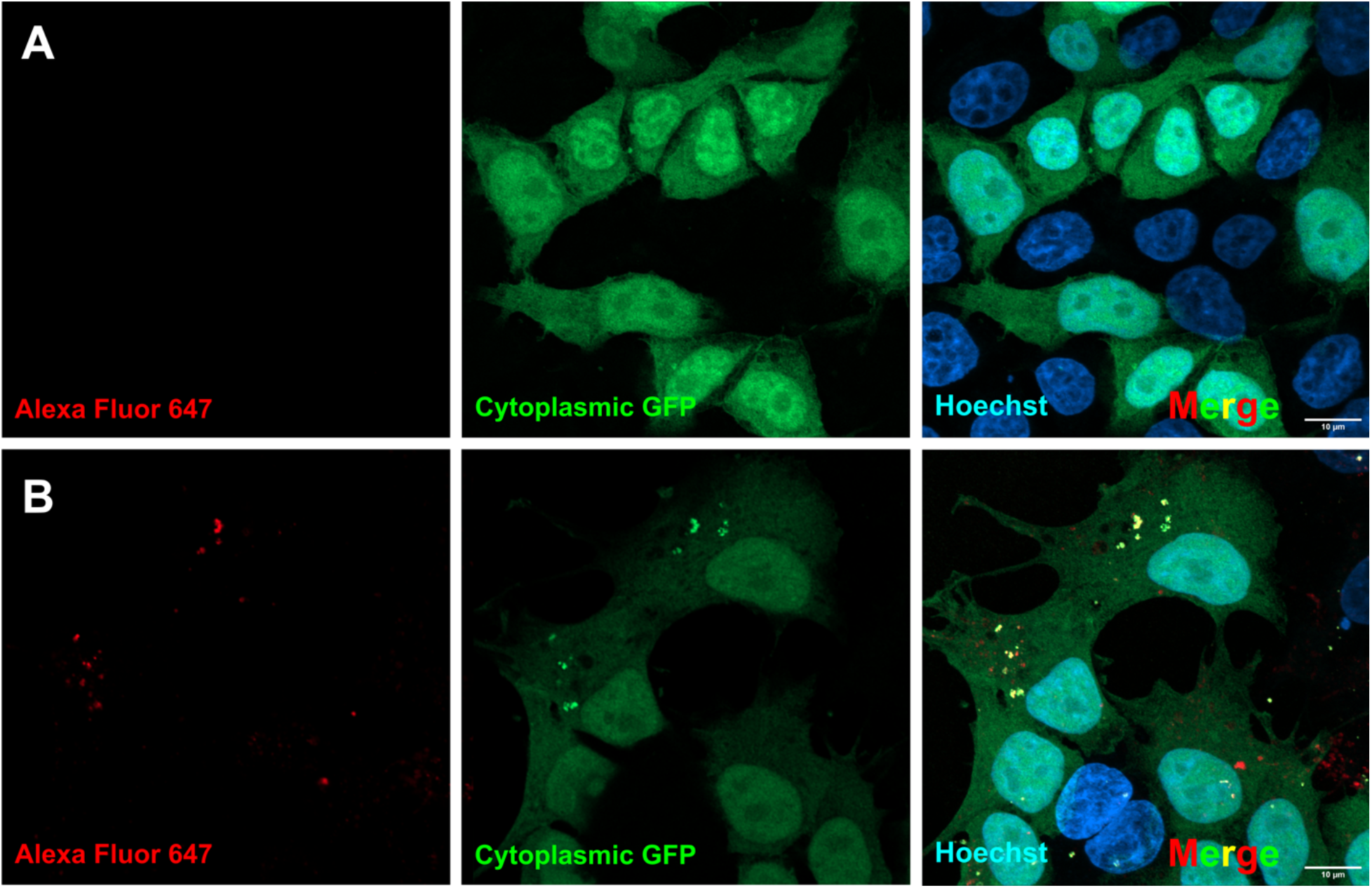
Visualization of colocalization of designed AF647-labeled protein cages and endogenous GFP. **(a)** Control experiment of cytoplasmic GFP without AF647-labeled protein cage. AF647, representing the fluorescently labeled protein cage, and cytoplasmic GFP can both be visualized using confocal microscopy. AF647 was not visualized by the confocal microscope due to the lack of protein cage. Without the protein cage, green puncta were unable to form and only diffuse cytoplasmic GFP can be visualized. **(b)** Colocalization of AF647-labeled protein cage and cytoplasmic GFP. Visualization of the AF647-labeled protein cage can be seen, resulting in the formation of green puncta. By merging the two images, colocalization of the protein cage and cytoplasmic GFP can be seen.

### Targeted imaging of specific proteins in live cells with designed protein cages

Our data suggests that fluorescently labeled protein cages can enter cells and target cytoplasmic proteins. We wanted to further investigate whether these cages can diffuse through different parts of the cell and have the potential to be targeted to specific cell compartments. To test this, we expressed a transmembrane mitochondrial protein, MFN1, fused to GFP. These proteins are fused to GFP at the N-terminus such that the GFP is presented as a cytoplasmic domain. Expression of MFN1-GFP in HEK-293T cells presents bright green puncta throughout the cytoplasm, in agreement with the localization and abundance of mitochondria present in HEK-293T cells. Next, we hypothesized that cages, once released into the cytoplasm, will diffuse until they recognize and bind the exposed GFP molecule and thus localize to MFN1 proteins on the surface of the mitochondria. Similarly, these cages are labeled with AF647 to enhance their visualization and localization in the cells.

After performing lipofection and introducing these labeled cages into the organelle-tagged cells, we found bright fluorescent puncta that colocalize between the green and far-red channels (**Fig. 6a-b**). This finding suggests that protein cages can be targeted to membrane-bound proteins belonging to specific cellular compartments.

**Figure 6.**
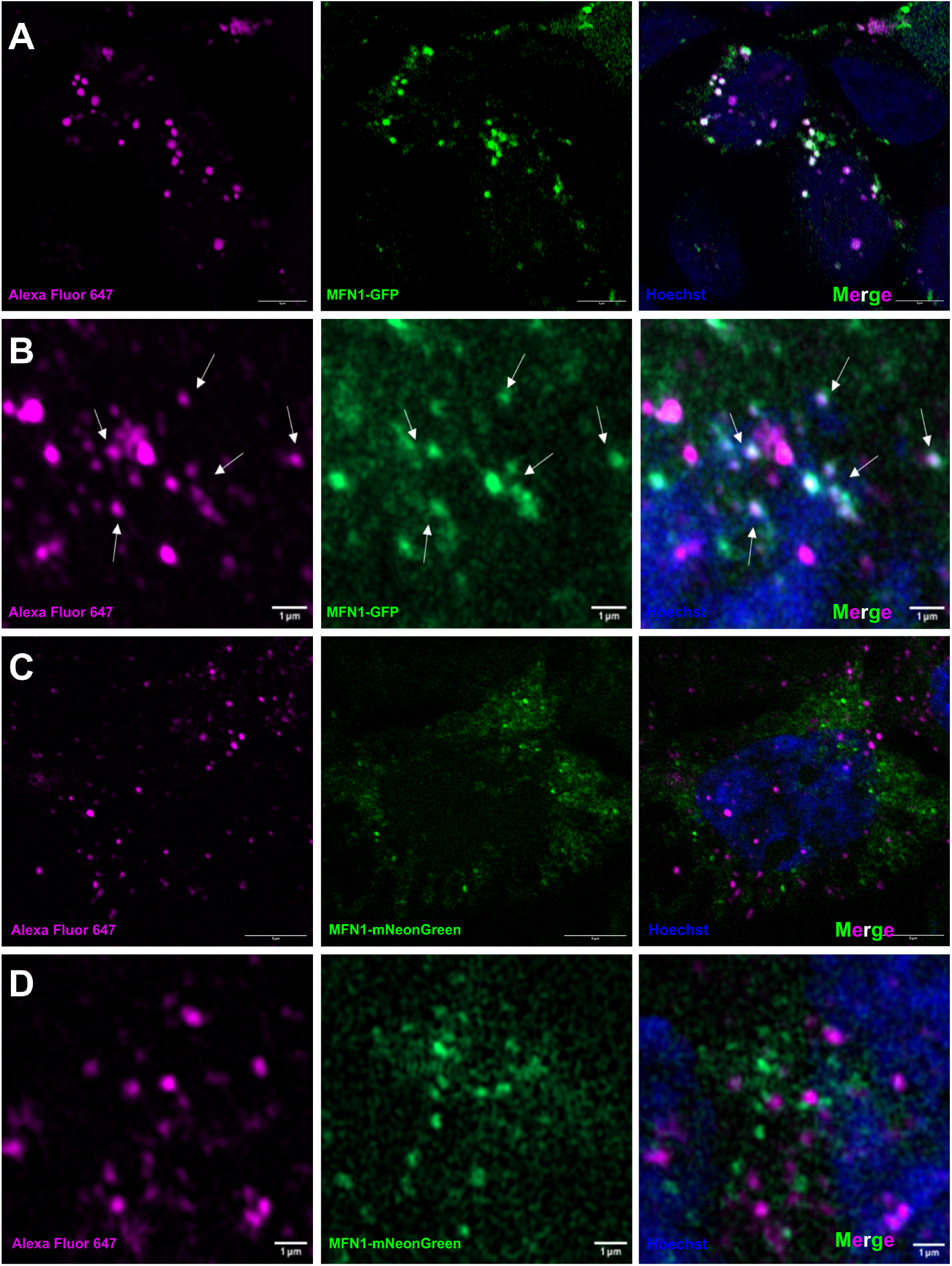
GFP-targeted cages are specific to GFP and do not display off-target binding to other fluorescent proteins. **(a-b)** Colocalization of AF647-labeled protein cages and the mitochondrial protein MFN1(labeled by GFP). By imaging AF647 emission (from the fluorescently labeled protein cage) and GFP emission (representing MFN1-GFP), and merging the two images, colocalization of the protein cage and GFP-tagged proteins can be visualized. **(c-d)** Lack of colocalization of AF647-labeled protein cages and MFN1-mNG. As the DARPins are unable to interact with mNG, the protein cages do not colocalize with MFN1.

To validate that the interaction is indeed specific to GFP and the DARPin domains on the protein cages, we also created HEK-293T cells expressing mNeonGreen (mNG) fused to MFN1. mNeonGreen is comparable in size and spectral properties to GFP but has almost no sequence similarity so we did not expect it to interact with the DARPins on the protein cages.

We found that, like MFN1-GFP, cells expressing MFN1-mNG also exhibit bright, green puncta reflective of mitochondrial distribution in the cell. However, when we transfected protein cages into HEK-293T cells expressing mNG-tagged MFN1, colocalization between the MFN1-mNG and the AF647 was not observed, indicating that cages labeled with the non-cognate DARPin (against GFP instead of mNG) were not targeted to the MFN1 protein or mitochondria (**Fig. 6c-d)**. Furthermore, the similar spectral properties of mNeonGreen and GFP provide a useful control to rule out autofluorescence or crosstalk between the far-red and green fluorescent channels as a complicating factor.

## DISCUSSION AND CONCLUSION

The central aim of this work was to develop innovative protein-based fiducial markers for cell imaging, with potential applications in molecular, cell, and structural biology research. We established that our large, designed, polyvalent protein cages could be internalized within HEK-293T cells and be readily visualized by fluorescence microscopy. The inclusion of DARPins in our protein cage design significantly enhances their modularity, allowing them to be targeted to specific cellular proteins or organelles. Using the mitochondrial transmembrane protein MFN1 as a first example, we showed that the location of this protein in the cell could be discerned. In this case study, the protein of interest (MFN1) was fused to GFP so that the initial protein cage construct (bearing an anti-GFP DARPin) could be used directly. For broader applications, including where genetic modification of a protein of interest is not possible or convenient, a suitably selected DARPin would be fused to the protein cage to direct it to the cellular protein of interest, in its native form. The prospective avenues for further development are myriad. Future research will focus on broader applications in mammalian cells along with more advanced imaging readouts in combination with cryo-CLEM.

The targeted binding of our designed protein cages to specific proteins inside cells also has potential applications in drug delivery. By modifying the protein cages to carry therapeutic cargo, such as small molecules, peptides, or nucleic acids, they could potentially be used to specifically deliver drugs to cells or subcellular compartments presenting a target protein, with the cargo being released upon internalization of the protein cage. The targeted binding of the protein cages could improve drug delivery efficiency and specificity, while reducing off-target effects and toxicity. In these initial experiments, the protein cages did not exhibit cell toxicity, making them potentially suitable for future *in vivo* explorations.

## MATERIALS AND METHODS

### Protein Production

The protein cage and the superfolder GFP V206A (sfGFP) were previously described ^19,26^. The proteins were expressed overnight at 18°C in Terrific Broth (Thermo Scientific) and 0.4% v/v glycerol with 0.5 mM isopropyl ß-D-1-thiogalactopyranoside (IPTG) induction at an O.D.600nm of 0.6-1.0. The cells were harvested by centrifugation at 4,000 xg for 20 minutes at 4°C.

For purification, cell pellets were resuspended in an amine-free extraction buffer [10 mM KCl, 220 mM NaCl, 50 mM NaHCO3, 20 mM Imidazole pH 8.0] in addition to benzonase nuclease (EMD Millipore), 1 mM PMSF, 1X EDTA-free protease inhibitor cocktail (Thermo Scientific), and 0.1% LDAO. The cells were lysed using an EmulsiFlex C3 homogenizer (Avestin), and the cell lysate was centrifuged at 20,000 xg for 20 minutes at 4°C. The remaining supernatant was centrifuged again at 5,000 x g for 20 minutes at 4°C and equilibrated for 1 hour in 3 mL of Ni-NTA resin (Qiagen) that was pre-equilibrated in the extraction buffer. Using a gravity column, the protein cage was eluted with elution buffer [10 mM KCl, 220 mM NaCl, 50 mM NaHCO3, 300 mM Imidazole pH 8.0] following a step elution model. Fractions containing the eluted protein cage were concentrated using an Amicon Ultra-15 100-kDa molecular weight cutoff centrifugal filter (MilliporeSigma), which were then purified with SEC. Using a Superose 6 Increase 10/300 GL column (Cytiva), the proteins were eluted with SEC buffer [5 mM KCl, 44 mM NaHCO3, 110 mM NaCl]. The chromatography fractions were analyzed using SDS-PAGE and negative stain electron microscopy to ensure proper assembly.

### Protein Labeling with Alexa Fluor 647 and sfGFP V206A

Protein cages were labeled with AF647 Antibody Labeling Kits (Invitrogen). Over 500 µg of protein cage was incubated per reaction for 1 hour at 25°C with periodic inversion, followed by an overnight incubation at 4°C ^42,43^. For sfGFP binding to the protein cage, a molar ratio of 2:1 of protein cage to sfGFP was mixed and incubated on ice for 5 minutes. Complex formation for both labeling systems was confirmed and isolated through SEC with a Superose 6 Increase 10/300 GL column.

### Negative stain electron microscopy

The concentration of a 3.5 μl sample of fresh SEC eluent was adjusted to ~100 μg/ml, applied to glow-discharged Formvar/Carbon 400 mesh Cu grids (Ted Pella Inc) for one minute and blotted to remove excess liquid. After a wash with filtered MilliQ water, the grid was stained with 2% uranyl acetate for one minute. Images were taken on a Tecnai T12 electron microscope.

### Cell transformation and lipofection

HEK-293T cells were grown under standard culture conditions at 5% CO2 with DMEM containing 10% fetal bovine serum. The cells were transfected with plasmids containing either GFP, MFN1-GFP, or MFN1-mNeonGreen, downstream of a CMV promoter to mark the cytoplasm or mitochondria, respectively.

Protein cages were packed into liposomes using Lipofectamine CRISPRmax Cas9 transfection reagents (Invitrogen). Protein cages were incubated with Lipofectamine CRISPRmax transfection reagents for 30 minutes in opti-MEM then added dropwise to cells. The lipofection mix was left on the cells for 4 hours, or 12 hours for MFN1 targeting experiments, before cells were fixed and processed for microscopy.

### Confocal imaging

HEK-293T cells were grown on 18 mm diameter circular coverslips in a 12-well dish in preparation for visualization. Coverslips were fixed with 4% PFA for 15 minutes then washed with PBS. Coverslips were then stained with Hoechst 33342 at a concentration of 1 μg/mL in PBS for 15 minutes then washed with PBS. Coverslips were then mounted with ProLong Glass Antifade Mountant (Thermo Scientific) on microscope slides. The mountant was left to cure overnight and confocal microscopy was performed the following day.

Super-resolution images were acquired on a Zeiss LSM 980 with Airyscan 2 using a 63x objective lens and appropriate lasers/filters for GFP, Hoechst, and AF647. Final images were analyzed using ImageJ ^44^.

## Acknowledgments

We thank Michael Stowell for suggestions and advice on cell targeting and imaging. We thank Nika Gladkov and Michael Sawaya for helping with the preparation of the cartoon picturing the protein cage. We thank Shannon Chau for helping with the preparation of the graphical abstract.

